# Multiplex genome editing by natural transformation (MuGENT) for synthetic biology in *Vibrio natriegens*

**DOI:** 10.1101/122655

**Authors:** Triana N. Dalia, Chelsea A. Hayes, Sergey Stolyar, Christopher J. Marx, James B. McKinlay, Ankur B. Dalia

**Author notes:** Author for correspondence: Ankur B. Dalia.

## Abstract

*Vibrio natriegens* has recently emerged as an alternative to *Escherichia coli* for molecular biology and biotechnology, but low-efficiency genetic tools hamper its development. Here, we uncover how to induce natural competence in *V. natriegens* and describe methods for multiplex genome editing by natural transformation (MuGENT). MuGENT promotes integration of multiple genome edits at high-efficiency on unprecedented timescales. Also, this method allows for generating highly complex mutant populations, which can be exploited for metabolic engineering efforts. As a proof-of-concept, we attempted to enhance production of the value added chemical poly-β-hydroxybutyrate (PHB) in *V. natriegens* by targeting the expression of nine genes involved in PHB biosynthesis via MuGENT. Within 1 week, we isolated edited strains that produced ~100 times more PHB than the parent isolate and ~3.3 times more than a rationally designed strain. Thus, the methods described here should extend the utility of this species for diverse academic and industrial applications.

---

*V. natriegens* is the fastest growing organism known, with a doubling time of <10 min^1,2^. With broad metabolic capabilities, lack of pathogenicity, and its rapid growth rate, it is an attractive alternative to *E. coli* for diverse molecular biology and biotechnology applications^3,4^. Methods for classical genetic techniques have been developed for V *natriegens*, but these are relatively laborious, require multiple steps, and must be used sequentially to generate multiple genome edits^3,4^. The challenges of these techniques contrast with the ease of genetics in *Vibrio* species that are naturally transformable. Competent *Vibrio* species can take up DNA from the environment and integrate it into their genome by homologous recombination; processes known as natural competence and natural transformation, respectively^5–8^. The inducing cue for natural transformation in competent *Vibrio* species is growth on the chitinous shells of crustacean zooplankton, which are commonly found in the aquatic environment where these microbes reside^5^. Chitin induces the expression of the competence regulator TfoX^9,10^. In fact, overexpression of TfoX obviates the need for chitin induction, allowing competent *Vibrio* species to be naturally transformed in rich media^5,9^.

As no reports of natural transformation existed for *V. natriegens*, we first sought to establish whether this was possible. Unlike naturally competent *V. cholerae*, incubation on chitin did not lead to detectable transformation in *V. natriegens* (data not shown). However, ectopic expression of TfoX (either the endogenous *tfoX* gene or one from *Vibrio cholerae)*on an IPTG-inducible plasmid (pMMB) supported high rates of natural transformation (**Fig. 1a**). This was tested using a linear PCR product that replaces the gene encoding the DNA endonuclease Dns with an antibiotic resistance (Ab^R^) marker. The *dns* locus was used as a target for transformation assays throughout this manuscript because loss of this gene does not impact growth or viability in rich medium. Under optimal conditions ~1–10% of the population had integrated the transforming DNA (tDNA), which matches the highest rates of transformation observed among competent species^11^ (**Fig. 1a-c**). Natural transformation of *V. natriegens* required very little transforming DNA (tDNA) (highly efficient with even 1 ng / 10^8^ CFU) and was dependent on the length of homologous sequence surrounding the mutation (**Fig. 1b and c**). This method could also be used to introduce point mutations into *V. natriegens* (tested with tDNA containing an *rpsL* K43R Sm^R^ allele); however, this activity was partially suppressed by the mismatch repair system (**Fig. 1d**).

**Fig. 1.**
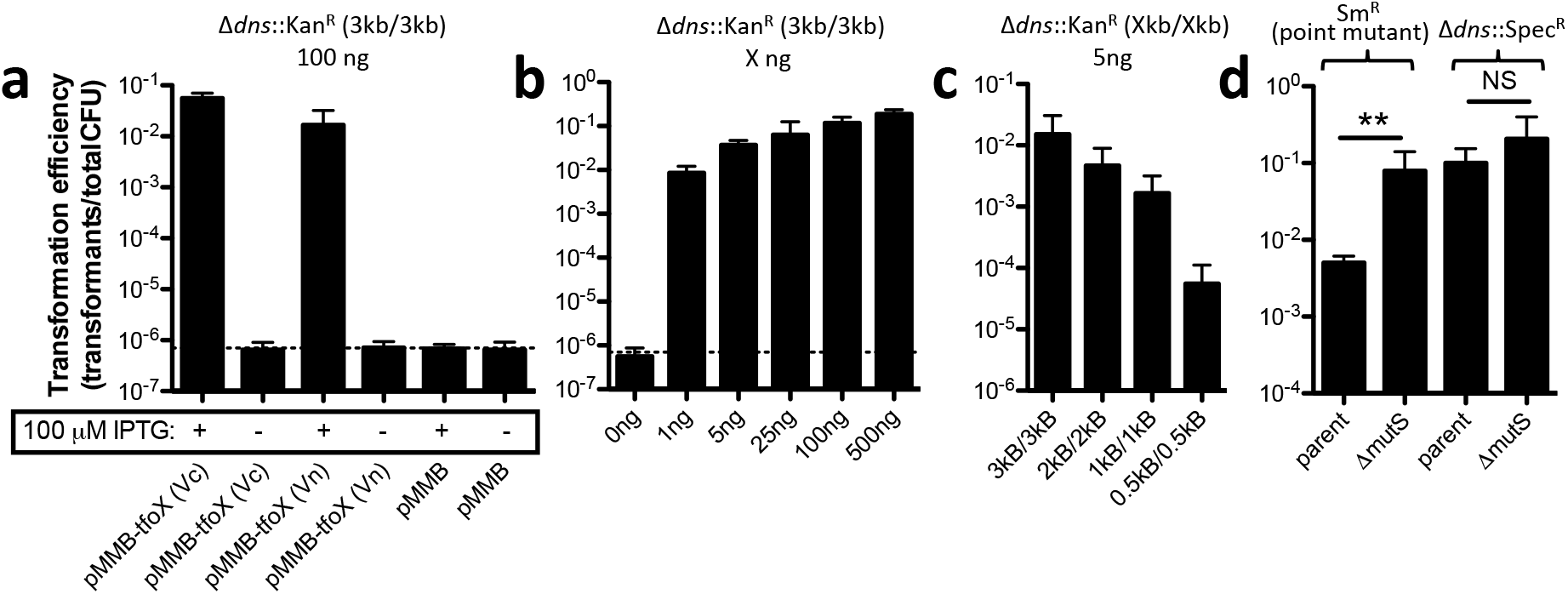
*Natural transformation of* V. natriegens. (**a-d**) Transformation assays of *V. natriegens*. (**a**) *V. natriegens* strains containing a pMMB empty vector or pMMB with the *tfoX* gene from either *V. natriegens* (Vn) or *V. cholerae* (Vc) were transformed with 100 ng of a Δ*dns*::Kan^R^ tDNA containing 3 kb flanks of homology on both sides of the mutation (i.e. 3 kb/3 kb). Transformation assay of *V. natriegens pMMB-tfoX* (Vc) with (**b**) the indicated concentration of Δ*dns*::Kan^R^ (3 kb/3 kb) tDNA or (**c**) 5 ng of Δ*dns*::Kan^R^ tDNA containing the indicated amount of homology on each side of the mutation. (**d**) Transformation assay in the indicated strain backgrounds with 5 ng of *rpsL* K43R Sm^R^ (3 kb/3 kb) or Δ*dns*::Spec^R^ (3 kb/3 kb) tDNA as indicated. All strains in **d** harbor Δ*dns*::Kan^R^ mutations and pMMB-tfoX (Vc). All data are shown as the mean ± SD and are the result of at least 4 independent biological replicates. ** = *p*<0.01.

Having demonstrated *V. natriegens* is naturally competent, we sought to determine if we could use natural transformation to perform scarless multiplex genome editing by natural transformation (MuGENT)^12^. MuGENT operates under the premise that under competence inducing conditions, only a subpopulation of cells is transformable. Those cells that can be transformed, however, have the capacity to take up and integrate multiple tDNAs^12,13^. Thus, during MuGENT, cells are incubated with two types of linear tDNA; (1) a selected product that introduces an antibiotic resistance marker into the genome and (2) unselected products that introduce scarless genome edits of interest at one or more loci.

We first tested the ability of MuGENT to introduce a single unmarked genome edit (also known as cotransformation). To facilitate measurement of cotransformation, we noted this species forms opaque colonies on agar plates (**Fig. 2a**), which could be due to the production of a capsular polysaccharide. Consistent with this, inactivating a homolog of the essential capsule biosynthesis gene *wbfF^14^* resulted in the formation of transparent colonies on agar plates and loss of expression of a high molecular weight polysaccharide (**Fig. 2a and 2b**). Thus, to test cotransformation we used an unselected product to replace ~500 bp of the 5’ end of the *wbfF* gene with a premature stop codon and scored cotransformation via colony morphology (opaque vs. transparent) on agar plates (**Fig. 3a**). We found that cotransformation was remarkably efficient in *V. natriegens* (up to ~80%), even with low amounts (~25–50 ng / 10^8^ CFU) of the unselected product (**Fig. 3b**). Also, cotransformation with 1 kb flanks on the unselected product was possible, but at ~6-fold lower frequencies than with 3 kb flanks (**Fig. 3c**).

**Fig. 2.**
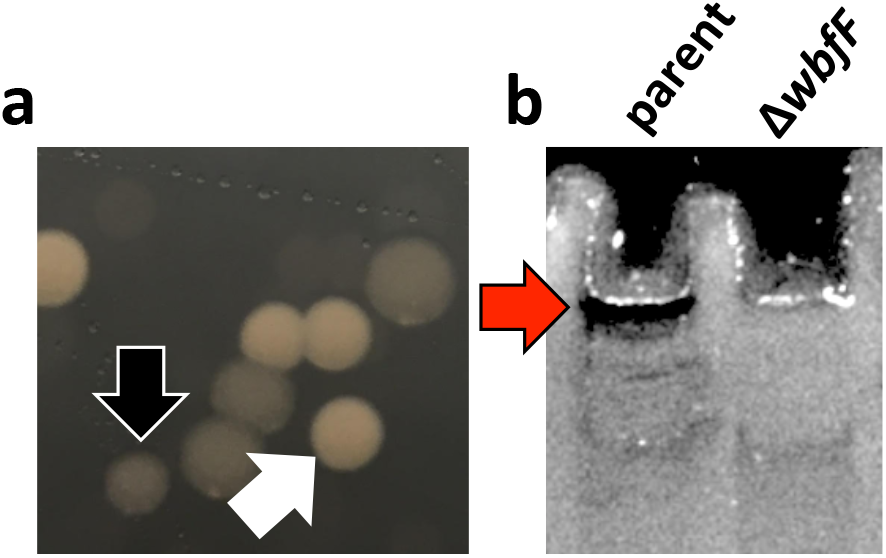
V. natriegens *produces a WbfF-dependent capsular polysaccharide*. (**a**) Colony morphologies of parent (white arrow) and Δ*wbfF* (black arrow) strains, which demonstrate the phenotypes screened for in cotransformation assays. (**b**) Cell lysates of the indicated strains were run on a 4-12% SDS PAGE gel and stained with the carbohydrate stain Alcian blue. The presence of a high molecular weight polysaccharide in the parent is indicated by a red arrow.

We next tested the full multiplex genome editing capacity of MuGENT to simultaneously cotransform multiple scarless genome edits into the genome in a single step^12,15^. Since there is no selection for integration of the unselected genome edits *in cis* during MuGENT, output populations are highly heterogeneous and individual mutants contain any number and combination of the multiplexed genome edits. Also, this process can be carried out in multiple iterative cycles to further increase the complexity of genome edits in the population (**Fig. 3d**)^12^.

**Fig. 3.**
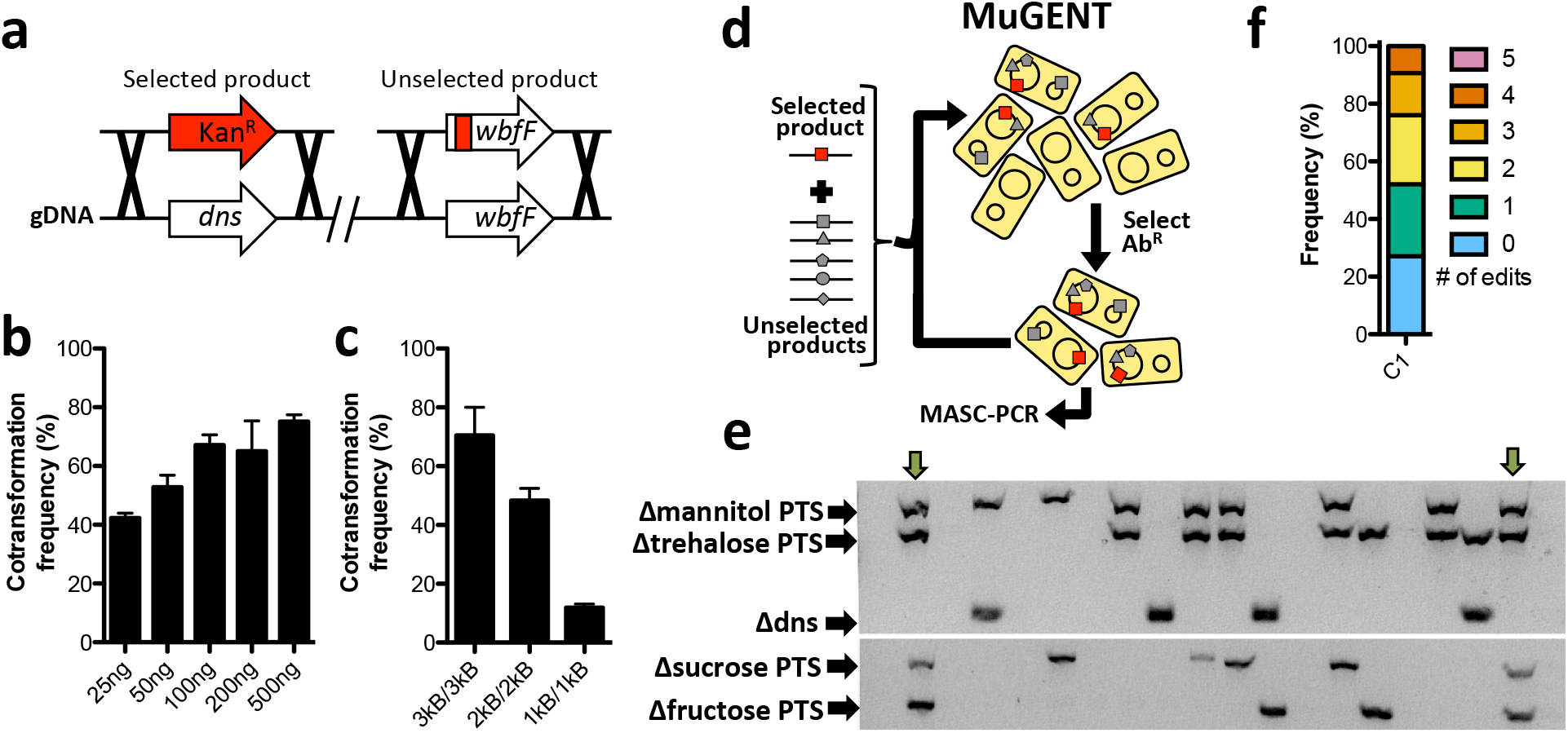
*Cotransformation is highly efficient in* V. natriegens. (**a**) Cotransformation was tested using a Δdns::Kan^R^ (3 kb/3 kb) selected product and an unselected product that deleted ~500 bp of the 5’ end of *wbfF* gene. Cotransformation assays were performed using 50ng of the Δdns::Kan^R^ (3 kb/3 kb) selected product and (**b**) the indicated amount of the *ΔwbfF* (3 kb/3 kb) unselected product or (**c**) 200 ng of *ΔwbfF* unselected products containing the indicated length of homology on each side of the mutation. Data in **b** and **c** are from at least four independent biological replicates and shown as the mean ± SD. (**d**) Schematic of MuGENT. The selected product is indicated by a red box, while multiple unselected genome edits are depicted by distinct gray shapes. Since there is no selection for genome edits *in cis*, output mutants can have any number and combination of the unselected genome edits. Circles inside cells represent the two circular chromosomes of *V. natriegens*. (**e** and **f**) MuGENT was performed with 5 unselected genome edits. The selected product was ΔwbfF::Kan^R^, while the unselected products targeted four carbohydrate transporters and *dns* for inactivation by replacing ~500 bp of the 5’ end of each gene with a premature stop codon. (**e**) A representative MASC-PCR gel of 24 colonies from the edited population. The targets of each genome edit are indicated on the left and the presence of a band indicates integration of the indicated genome edit. Strains containing 4 genome edits are indicated by the green arrows. (**f**) Distribution of genome edits in the population determined by MASC-PCR analysis of 48 random mutants.

As an initial test of multiplex genome editing, we targeted 5 genes whose mutagenesis was considered unlikely to affect viability or growth in LB. These targets included four carbohydrate transporters (specific for mannitol, fructose, sucrose, and trehalose – all of which are absent in LB) and the *dns* gene. All genes were targeted for inactivation by replacing ~500 bp of the 5’ end of each gene with a premature stop codon. Integration of genome edits was determined by multiplex allele-specific colony PCR (MASC-PCR)^16^ (**Fig. 3e**). Following one cycle of MuGENT, we found that ~70% of the population contained at least 1 genome edit, with ~25% of the population containing 3-4 genome edits (**Fig. 3f**). A quadruple mutant from this experiment was isolated and whole genome sequencing of this strain did not reveal any off-target mutations. Thus, MuGENT rapidly generated *V. natriegens* strains with multiple large (0.5 kb) scarless genome edits at high-efficiency without off-target effects, and can be used to make highly complex mutant populations.

As a second demonstration of multiplex genome editing, we demonstrated its utility in metabolic engineering by attempting to rapidly enhance production of a value-added chemical in *V. natriegens*. This species naturally accumulates low levels of the bioplastic precursor poly-β-hydroxybutyrate (PHB) as a storage polymer^17^. PHB is derived from the condensation and subsequent NADPH-dependent reduction of acetyl-CoA precursors^18^. Thus, for our targets, we tuned the expression (swap out *Pnative* for IPTG-inducible Ptac) or inactivated genes that we hypothesized would affect NADPH and/or acetyl-CoA availability. The targets for promoter swaps were the PHB synthesis operon (*phaBAC*), NAD kinase (*nadK*), and two transhydrogenases (*pntAB* and *udhA*), while targets for inactivation were phosphoglucose isomerase (*pgi*), citrate synthase (gltA), phosphotransacetylase (*pta*), isocitrate lyase (aceA), and lactate dehydrogenase (*ldhA*) (**Fig. 4a**). Thus, there were 512 possible combinations for these 9 genome edits. We performed multiple cycles of MuGENT to introduce these genome edits into a competent population of *V. natriegens*. At each cycle, the selected product was designed to swap the Ab^R^ marker at the *dns* locus to maintain coselection at each step. Following four cycles of MuGENT, which took just 5 days to perform, ~50% of the population had 3 or more genome edits and ~10% contained 5+ genome edits (**Fig. 4b**). To select mutants with increased PHB production, we then plated this output population onto media containing Nile red, which stains PHB granules^19^. Nile red fluorescence on these plates was highly heterogeneous, suggesting that some genotypes produced more PHB than the parent isolate (**Fig. 4c**). A number of highly fluorescent colonies were picked and the genotypes determined by MASC-PCR. Also, PHB in these select strains was directly measured by HPLC. Cumulatively, these analyses rapidly revealed genotypes that produced ~100-fold more PHB than the parent and ~3.3-fold more than a strain with just the *P_tac^-^_phaBAC* mutation (**Fig. 4d**). Overexpression of the *phaBAC* locus is a commonly used rational approach for enhancing PHB production ^18,20^. Thus, this result demonstrates that MuGENT can allow for rapid isolation of genotypes associated with enhanced phenotypes (e.g. enhanced PHB production) compared to rationally engineered strains (e.g. a *P_tac^-^_phaBAC* mutant) without prior knowledge of effective combinations of individual mutations.

**Fig. 4.**
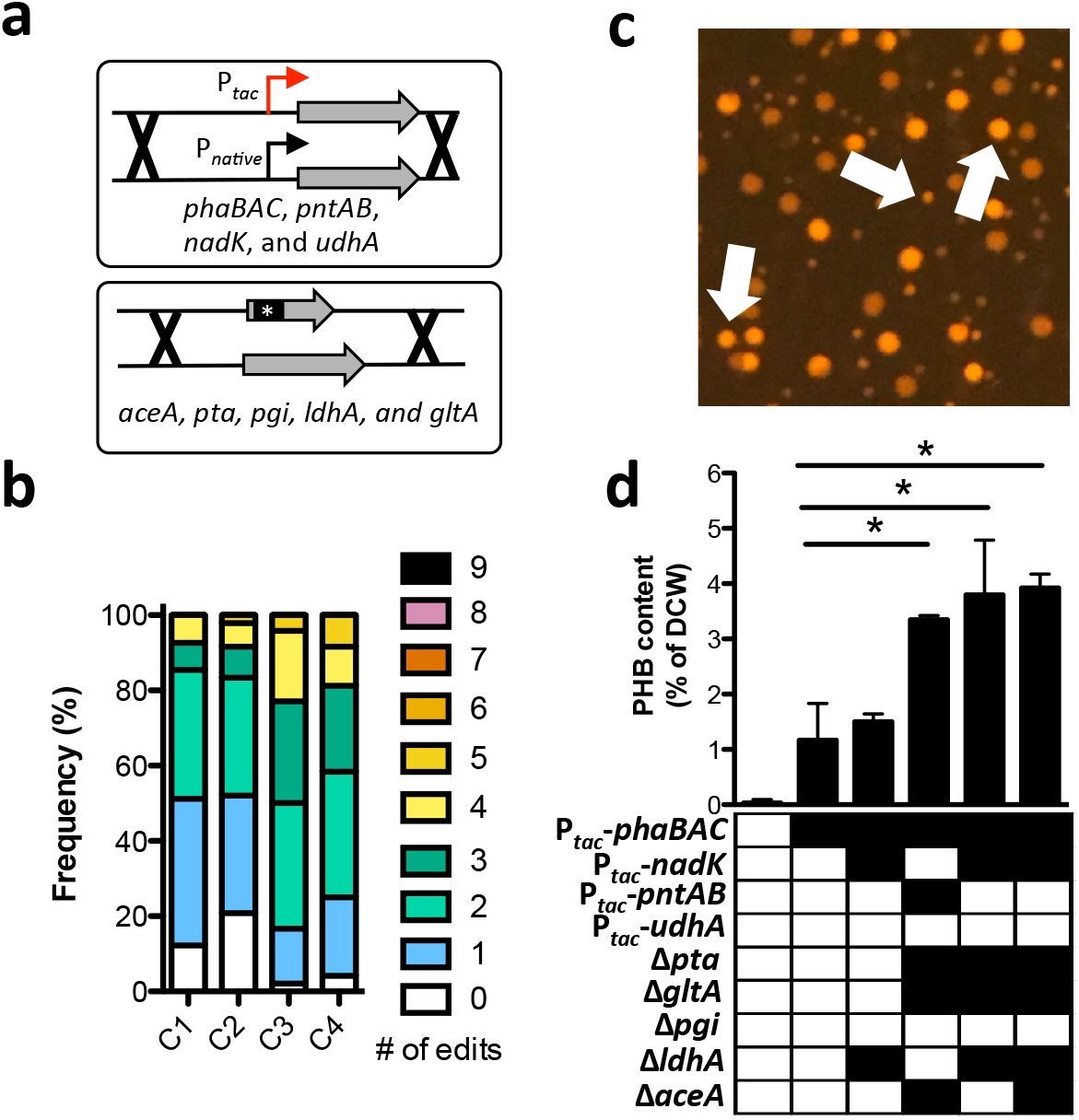
*MuGENT rapidly enhances PHB production in* V. natriegens. (**a**) The indicated targets were subjected to either a promoter swap (top) or inactivation by replacing ~500bp of the 5’ end of each gene with a short sequence to introduce a premature stop codon (bottom). (**b**) Distribution of the 9 genome edits in a population of cells following four cycles of MuGENT. (**c**) Representative image of the mutant pool generated in b plated on Nile red containing plates, which stain PHB granules. White arrows indicate colonies with increased fluorescence intensity compared to the parent. (**d**) PHB content of select MuGENT optimized strains is shown as the % of dry cell weight (DCW). The genotype of each mutant is shown below each bar where a filled box indicates the presence of the genome edit indicated on the left. Data are shown as the mean ± SD and are from at least 2 independent biological replicates. * = *p*<0.05.

While many methods for multiplex genome editing in bacterial systems have been described^21^, many of these are limited to small changes such as SNPs. MuGENT, on the other hand, can efficiently swap, insert, or remove whole promoters or coding sequences as demonstrated above. Furthermore, one of the major limitations to other multiplex genome editing methods is that mutagenesis must be performed in strains lacking DNA repair pathways to allow for high-efficiency integration of genome edits, which results in a large number of off-target mutations^16,21^. MuGENT in *V. natriegens* is performed in DNA repair sufficient backgrounds, thus, little to no off target mutations are introduced during the procedure as indicated above. Also, unlike other multiplex editing approaches, MuGENT requires no specialized equipment and, thus, has the potential to make multiplex genome editing commonplace.

In conclusion, this study demonstrates that MuGENT is a rapid, efficient, and simple tool for engineering the *V. natriegens* genome. This microbe is already being developed as an alternative to *E. coli*, and we believe that the ease and speed of MuGENT will extend the use of *V. natriegens* as a novel chassis for diverse molecular biology and biotechnology applications.

## METHODS

### Bacterial strains and culture conditions

The parent *V. natriegens* strain used throughout this study was a spontaneous rifampicin-resistant derivative of ATCC 14048^2^. For a list of all strains used / generated in this study, see **Table S1**. Strains were routinely grown in LB+v2 salts (LBv2)^3^, which is LB Miller broth (BD) supplemented with 200 mM NaCl, 23.14 mM MgCl2, and 4.2 mM KCl. LBv2 was supplemented with 100 μM IPTG, 50 μg/mL kanamycin (Kan), 200 μg/mL spectinomycin (Spec), 100 μg/mL rifampicin (Rif), 100 μg/mL streptomycin (Sm), or 100 μg/mL carbenicillin (Carb) as appropriate.

### Generation of mutant strains and constructs

Mutant constructs were generated by splicing-by-overlap extension (SOE) PCR exactly as previously described^22^. Briefly, for three-piece mutant constructs (i.e. for constructs where a gene of interest is replaced with an Ab^R^ cassette or where the native promoter is swapped for a Ptac promoter) segments were designated UP, MIDDLE, and DOWN and correspond to: (1) UP = the upstream region of homology amplified with F1 and R1 primers, (2) DOWN = the downstream region of homology amplified with F2 and R2 primers, and (3) MIDDLE = the Ab^R^ marker or promoter swap fragment. For two-piece mutant constructs (i.e. for constructs where ~501 bp of the 5’ end of a gene is replaced with a stop codon), the mutation of interest is incorporated into the R1 and F2 primers used to amplify the upstream and downstream regions of homology, respectively. Gel purified segments were then mixed in equal ratios and used as template for a SOE PCR reaction with the F1 and R2 primers. All mutant constructs were made using Phusion polymerase. These were introduced into the *V. natriegens* genome via natural transformation as described below. All primers used to generate mutant constructs are listed in Table S2.

### Natural transformation / MuGENT assays

Strains harboring pMMB-tfoX (Vn *tfoX* or Vc *tfoX*) were induced to competence by growing overnight (12-18 hours) in LBv2+100 μg/mL carbenicillin+100 μM IPTG in a rollerdrum at 30°C. Then, ~10^8^ CFUs of this overnight culture (~3.5 μL) were diluted directly into 350 μL of instant ocean medium (28 g/L; Aquarium Systems Inc.) supplemented with 100 μM IPTG. Transforming DNA (tDNA) was then added as indicated, and reactions were incubated statically at 30°C for 5 hours. Next, 1 mL of LBv2 was added and reactions were outgrown at 30°C with shaking (250 rpm) for ~1-2 hrs. Then, reactions were plated for quantitative culture onto media to select for integration of tDNA (i.e. LB+drug = transformants) and onto nonselective media (i.e. plain LB = total viable counts). Transformation efficiency is shown as: transformants / total viable counts.

For MuGENT, transformation assays were conducted exactly as described above. Unless otherwise specified, ~50 ng of the selected product was incubated with cells along with ~200 ng of each unselected product. After outgrowth, 1/10^th^ of the reaction was removed and plated for MASC-PCR analysis (described below). If multiple cycles of MuGENT were performed, the rest of the reaction was grown overnight in LBv2 supplemented with 100 μM IPTG, 100μg/mL carbenicillin (to maintain *p MMB-tfoX*), and the antibiotic to select for integration of the selected product. The following day, the population was then subjected to another round of MuGENT as described above using a selected product containing a different Ab^R^ marker to maintain coselection at each cycle.

Integration of genome edits was detected via MASC-PCR exactly as previously described^12,16^. Briefly, colonies were boiled in 50 μL of sterile water, vortexed, and then 2 μL were used as template in a 25 μL PCR reaction. PCR was conducted with Taq polymerase (SydLabs) using a modified 5X Taq buffer: 200 mM Tris pH 8.8, 100 mM KCl, 100 mM (NH4)2SO4, 30 mM MgSO4, and 1% Triton X-100. The total primer used in each MASC-PCR reaction (regardless of the number of multiplexed products being detected) was 1200 nM (i.e. for detection of 4 multiplexed genome edits, 300 nM of each genome edit-specific primer pair was used). The cycling conditions used were: 95°C 3 min; 26 × [95°C 40s, 58°C 30s, 72°C 3 min]; 72°C 3 min; 12°C hold. Reactions were then run on 2% agarose gels and imaged with GelGreen dye according to manufacturer’s instructions (Biotium). For a list of all primers used for MASC-PCR see **Table S2**.

### Alcian blue stained gels

To prepare cell lysates, ~10^9^ cells of the indicated *V. natriegens* strains were pelleted and then resuspended in 180 μL of Buffer ATL (Qiagen). Then, 20 μL of a 20 mg/mL proteinase K stock solution was added to each reaction and incubated at 56°C for 20 mins. Samples were then boiled in 2X SDS PAGE sample buffer and separated on 4-12% SDS PAGE gels. Gels were then stained with 0.1% Alcian Blue 8GX in 40% ethanol/3% acetic acid as previously described^23^. The gel was then destained in a 40% ethanol/3% acetic acid and imaged on a Biorad ChemiDoc MP Imaging system.

### Whole genome sequencing

Genomic DNA was prepped from strains and sequencing libraries were prepped via homopolymer-tail mediated ligation exactly as previously described^24^. Single-end 50 bp reads were collected on the Illumina platform. Then, data was analyzed for small indels and single nucleotide variants using CLC Genomics Workbench exactly as previously described^15,25^.

### Qualitative and quantitative assessment of PHB production

PHB was qualitatively assessed in MuGENT edited populations of *V. natriegens* by plating onto Nile red containing medium with excess glucose as a carbon source and 100 μM IPTG to induce P*_tac_*-containing genome edits = recipe per L: 28 g instant ocean, 2.5 g tryptone, 1 g yeast extract, 20 g glucose, 15 g agar, and 1 mg Nile red. Fluorescence of colonies was detected using a PrepOne Sapphire LED blue light base (475 nm ± 30 nm) and amber filter (530 nm long pass) (Embi Tec).

For quantitative assessment of PHB levels, the indicated strains were grown overnight in M9 minimal medium (BD) supplemented with 2 mM MgSO4, 100 μM CaCl2, 200 mM NaCl, 30 μM FeSO4, 100 μM IPTG, 1% tryptone, and 2% glucose. Approximately 8 × 10^9^ cells were then pelleted, resuspended with 50 μL water and transferred to pre-weighed glass screwcap tubes. Cell suspensions were dried for 5 h at 80°C and then the tubes were weighed again to determine dry cell weights. PHB was then hydrolyzed and extracted as crotonic acid by boiling the dried cells in 1 ml of pure sulfuric acid. Extracts were chilled on ice and diluted with 4 ml ice-cold water. Aliquots were further diluted 10-fold with water, centrifuged, filtered, and then crotonic acid was quantified by HPLC as described^26^.

## ACKNOWLEDGEMENTS

We would like to thank Tufts TUCF Genomics and the Indiana University CGB for assistance with whole genome sequencing of strains. This work was supported by US National Institutes of Health Grant AI118863 to ABD. JBM and ABD were also supported by the Indiana University College of Arts and Sciences.

## AUTHOR CONTRIBUTIONS

ABD conceived the study. TND, SS, CJM, JMB, and ABD designed experiments. TND, CAH, JMB, and ABD performed experiments. All authors played a role in writing and/or proofreading the manuscript.

## SUPPORTING INFORMATION

Supplementary protocol: Natural transformation / MuGENT in *V. natriegens*.Supplementary Tables S1-S2: strains and primers used in this study.

## Supplementary material for

### Contents

**Supplementary protocol** – *Natural transformation / MuGENT in V. natriegens*

**Table S1** – *Strains used in this study*

**Table S2** – *Primers used in this study*

## Supplementary protocol: Natural transformation / MuGENT in *V. natriegens*

### Materials

1. LBv2 = Mix 400 mL LB +32 mL sterile 5 M NaCl + 3.4 mL sterile 1 M KCl + 18.5 mL sterile 1 M MgCl2
2. 2XIO = 28 g/L of instant ocean sea salts (www.instantocean.com)
3. Transforming DNA (tDNA) –

a. selected product = mutant construct that has a selectable marker (antibiotic resistance cassette) that replaces the gene of interest or a neutral locus (for MuGENT). Homology on each side of the mutation can be as little as 0.5 kb. Homology of 3 kb on either side of the mutation results in the highest transformation efficiencies.
b. Unselected product (cotransformation / MuGENT) = mutant construct that lacks any selectable marker but has a mutation of interest (deletion, point mutation, promoter swap, etc.). Unselected products should have 3 kb of homology on each side of the mutation for the highest rates of cotransformation / MuGENT.

### Notes

1. Carbl00 = Carbenicillin 100 μg/mL
2. SAD1306 =*V. natriegens* Rif^R^ 14048 pMMB67EH-tfoX
3. The pMMB67EH-tfoX plasmid is very stable in *V. natriegens*, therefore, Carbenicillin (Carb) is not needed to maintain it throughout the transformation protocol.

### Procedure

1. Inoculate 3 mL of LBv2+Carb100+100 μM IPTG in a culture tube with SAD1306. Grow at 30°C in rollerdrum overnight (12-18 hours).
2. Next day, take culture out of incubator and measure OD_600_. Generally, our overnights are at an OD_600_ of between 7-10.
3. For each transformation reaction, take 3.5 μL of the overnight culture and dilute into 350 μL of 2XI0+100 μM IPTG (no Carb). Invert gently to mix. Be sure to also prep a “no DNA” control reaction.
4. Add tDNA to each reaction and invert gently to mix:

a. For a selected product (i.e. a product that has an antibiotic resistance marker) = ~5-50 ng yields thousands of colonies.
b. For cotransformation / MuGENT = use 50 ng of a selected product and ~200 ng of each unselected product
5. Incubate reactions at 30°C statically for 4-6 hours.
6. Next, add 1mL LBv2 (no drug) to each transformation reaction and outgrow at 30°C shaking for 1-2 hours
7. To determine the transformation efficiency:

a. Plate all reactions for quantitative culture on media to select for the transformants (i.e. on antibiotic plates that select for integration of selected product) and on plates without any drug to determine the total CFU in the culture.
b. Transformation efficiency = transformants CFU / total CFU
8. For cotransformation / MuGENT:

a. Plate onto media to select for integration of the selected product.
b. Pick single colonies and screen by MASC-PCR to identify clones with the desired genome edits.
c. To perform a subsequent round of MuGENT, ~200 μL of the outgrown transformation can be inoculated into 3 mL of LBv2+Carb100+100 μM IPTG+the antibiotic that selects for integration of the selected product (e.g. if Δdns::Kan^R^ was used in the first cycle of MuGENT, then 50 μg/mL Kan would be included in this overnight culture). Start at “step 2” of this procedure to perform the next cycle of MuGENT being sure to use a selected product with a distinct Ab^R^ marker.

## Supplementary Tables

**Table S1.**
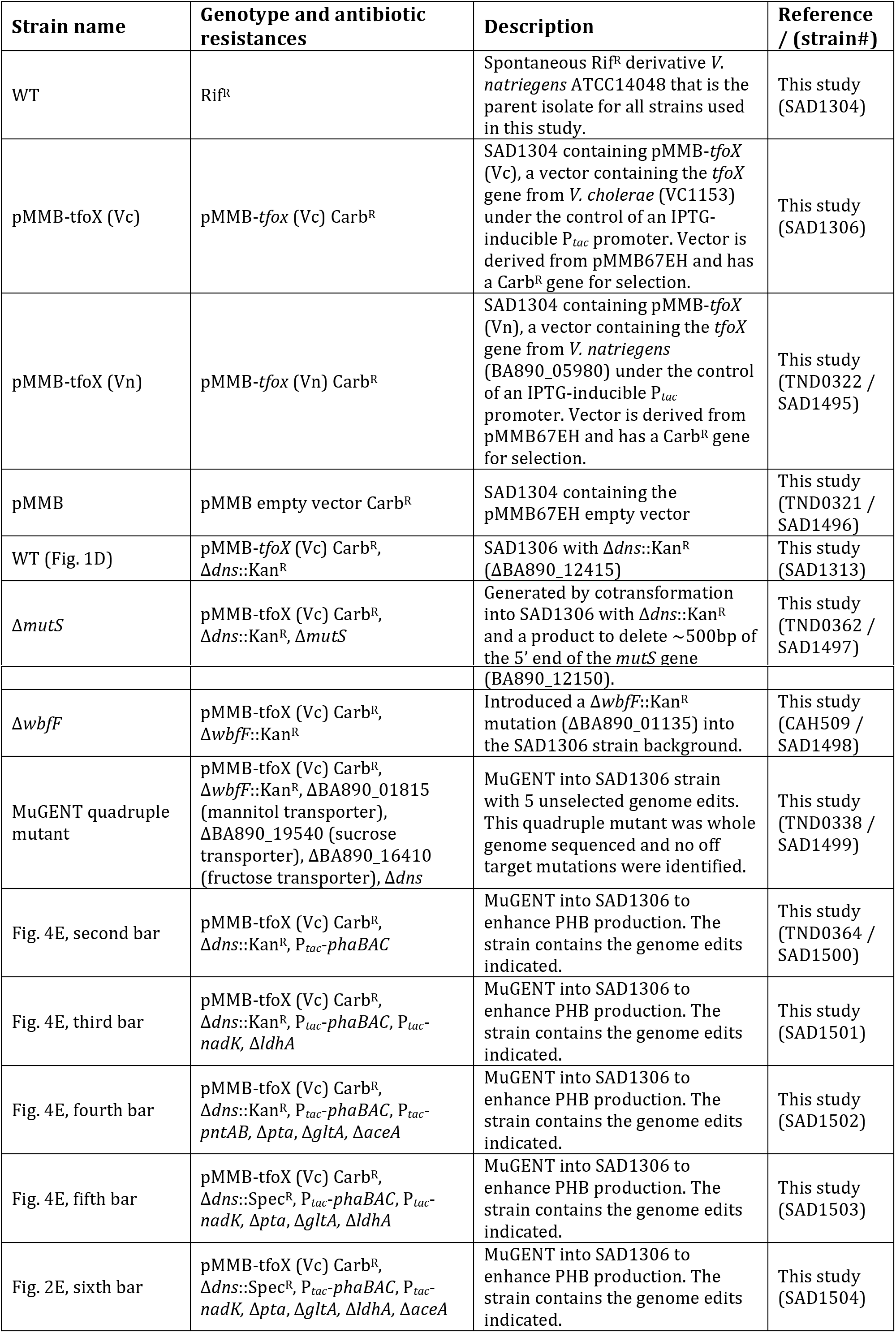
*Strains used in this study*

**Table S2.**
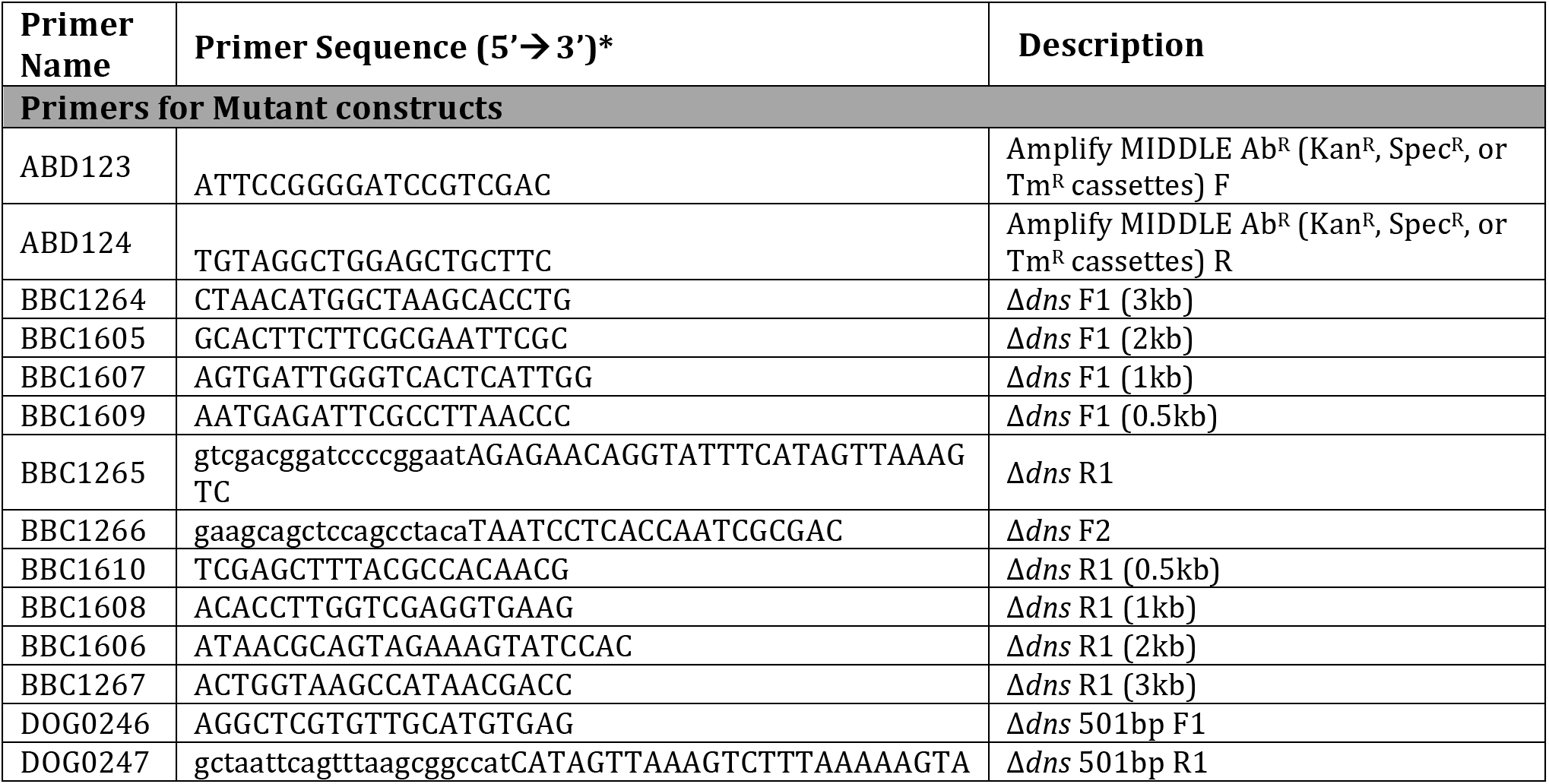
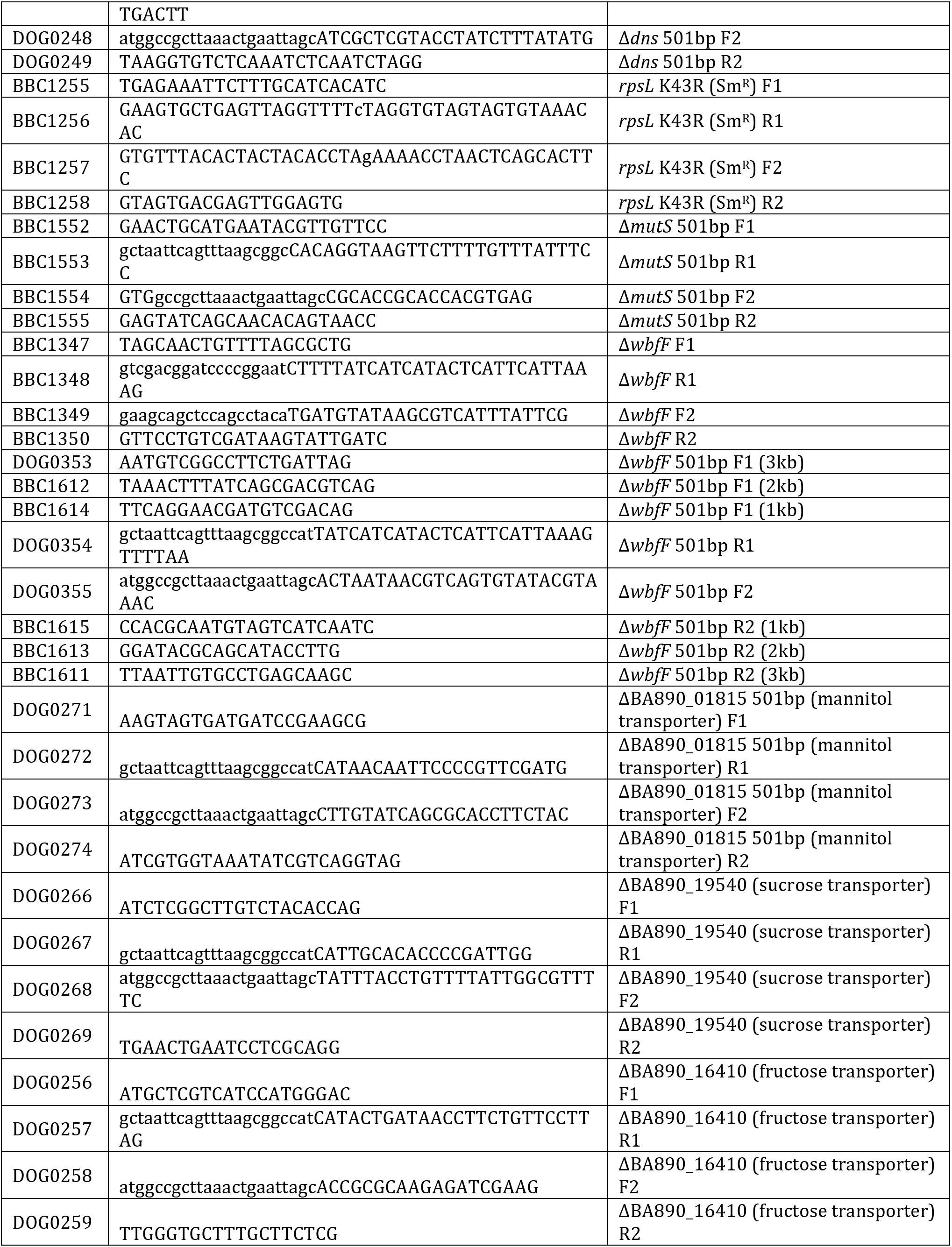
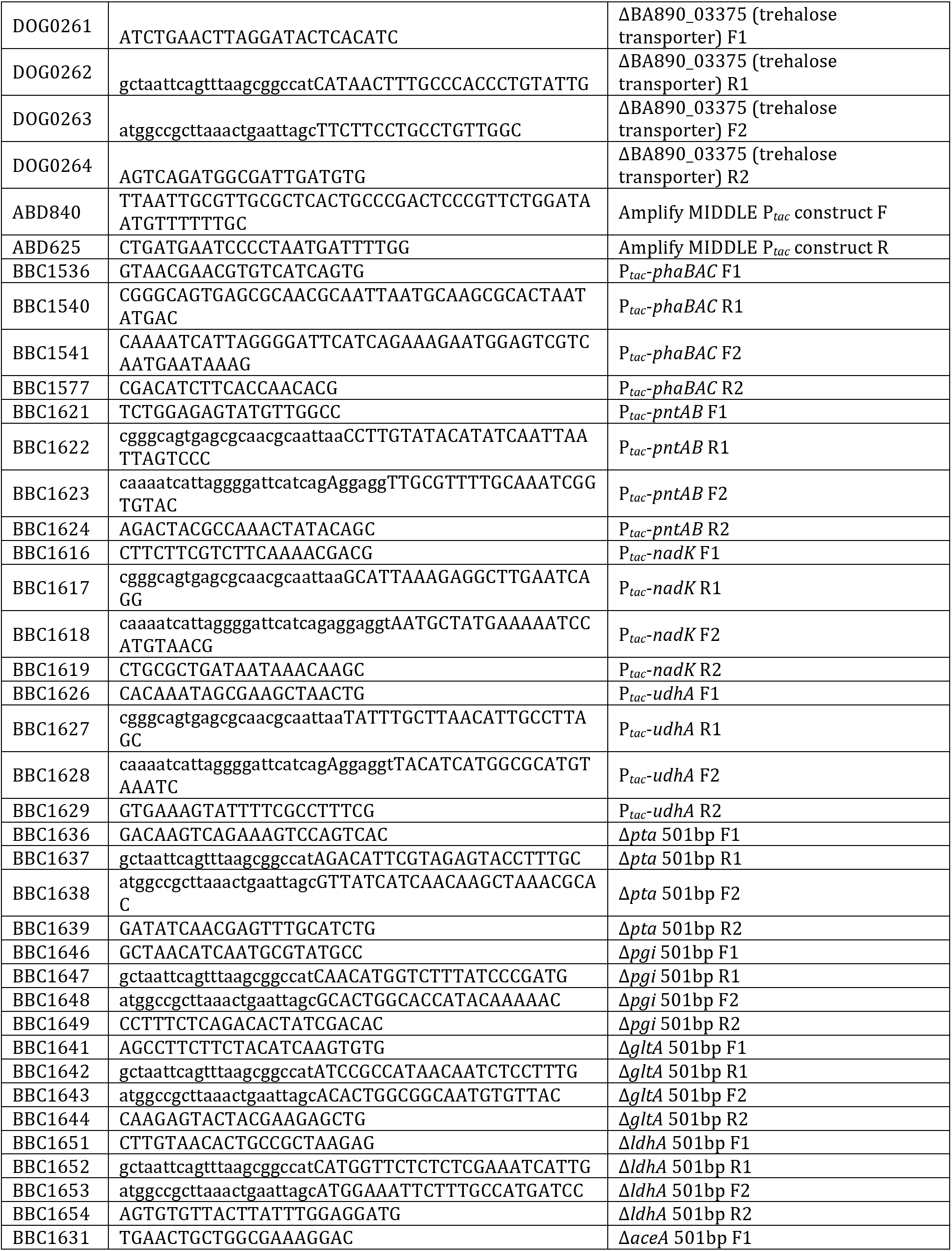
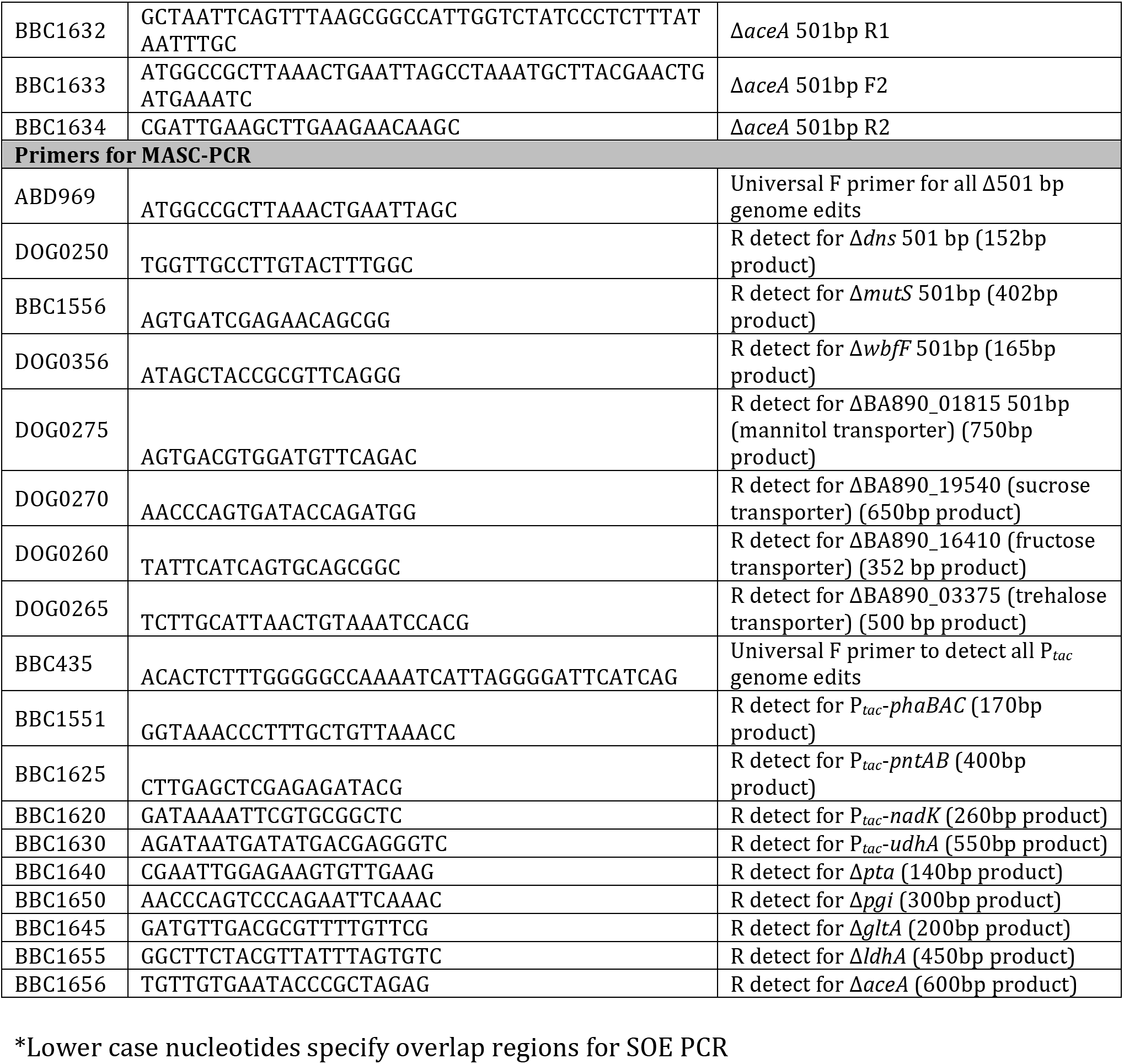
*Primers used in this study*

